# The Structure of ChAdOx1/AZD-1222 Reveals Interactions with CAR and PF4 with Implications for Vaccine-induced Immune Thrombotic Thrombocytopenia

**DOI:** 10.1101/2021.05.19.444882

**Authors:** Alexander T. Baker, Ryan J. Boyd, Daipayan Sarkar, John Vant, Alicia Teijeira Crespo, Kasim Waraich, Chloe D. Truong, Emily Bates, Eric Wilson, Chun Kit Chan, Magdalena Lipka-Lloyd, Petra Fromme, Marius Bolni Nagalo, Meike Heurich, Dewight Williams, Po-Lin Chiu, Pierre J. Rizkallah, Alan L. Parker, Abhishek Singharoy, Mitesh J. Borad

## Abstract

Adenovirus derived vectors, based on chimpanzee adenovirus Y25 (ChAdOx1) and human adenovirus type 26 are proving critical in combatting the 2019 SARS-CoV-2 pandemic. Following emergency use authorisation, scale up in vaccine administration has inevitably revealed vaccine related adverse effects; too rare to observe even in large Phase-III clinical trials. These include vaccine-induced thrombotic thrombocytopenia (VITT), an ultra-rare adverse event in which patients develop life-threatening blood clots 5-24 days following vaccination.

To investigate vector-host interactions of ChAdOx1 underpinning VITT we solved the structure of the ChAdOx1 capsid by CryoEM, and the structure of the primary receptor tropism determining fiber-knob protein by crystallography. These structural insights have enabled us to unravel key protein interactions involved in ChAdOx1 cell entry and a possible means by which it may generate misplaced immunity to platelet factor 4 (PF4), a protein involved in coagulation.

We use in vitro cell binding assays to show that the fiber-knob protein uses coxsackie and adenovirus receptor (CAR) as a high affinity binding partner, while it does not form a stable interface with CD46. Computational simulations identified a putative mechanism by which the ChAdOx1 capsid interacts with PF4 by binding in the spaces between hexon proteins, with downstream implications for the causes of VITT.

**Summary:** We present the structure of the ChAdOx1 viral vector, derived from chimpanzee adenovirus Y25 at 4.2Å resolution^1^. ChAdOx1 is in global use in the AstraZeneca vaccine, ChAdOx1 nCoV-19/AZD-1222, to combat the SARS-CoV-2 coronavirus pandemic. Recently observed, rare, adverse events make detailed mechanistic understanding of this vector key to informing proper treatment of affected patients and the development of safer viral vectors.

Here, we determine a primary mechanism ChAdOx1 uses to attach to cells is coxsackie and adenovirus receptor (CAR), a protein which is identical in humans and chimpanzees. We demonstrate the vector does not form a stable CD46 interaction, a common species B adenovirus receptor, via its primary attachment protein.

Further, we reveal the surface of the ChAdOx1 viral capsid has a strong electronegative potential. Molecular simulations suggest this charge, together with shape complementarity, are a mechanism by which an oppositely charged protein, platelet factor 4 (PF4) may bind the vector surface. PF4 is a key protein involved in the formation of blood clots^2^, and the target of auto-antibodies in heparin-induced immune thrombotic thrombocytopenia (HITT)^3^, an adverse reaction to heparin therapy which presents similarly to vaccine-induced immune thrombotic thrombocytopenia (VITT), a rare complication of ChAdOx1 nCoV-19 vaccination^4–6^. We propose a mechanism in which the ChAdOx1-PF4 complex may stimulate the production of antibodies against PF4, leading to delayed blood clot formation, as observed in VITT.

## Introduction

The ChAdOx1 viral vector, adapted from chimpanzee adenovirus Y25 (ChAd-Y25), serves as the basis for the ChAdOx1 nCoV-19 vaccine, also called AZD-1222^1^. ChAdOx1 nCoV-19, distributed by AstraZeneca, induces robust immunity against the severe acute respiratory coronavirus 2 (SARS-CoV-2), protecting against severe symptoms requiring hospitalisation, in 100% of recipients, and infection of any severity, in approximately 70%^7,8^. However, it appears a minority of recipients develop a life-threatening clotting disorder which presents similarly to heparin induced thrombotic thrombocytopenia (HITT) as a result of receiving this vaccine^4,6^. Similar observations have been made in recipients of the Janssen designed Ad26.COV2.S vaccine, distributed by Johnson & Johnson, which is derived from the species D human adenovirus type 26 (HAdV-D26)^5,6^. The mechanism which results in this condition, termed vaccine-induced immune thrombotic thrombocytopenia (VITT), is unknown, although recent reports have highlighted a possible role for platelet factor 4 (PF4)^9^.

Detailed mechanistic understanding of the host-vector interactions of adenovirus derived vectors has facilitated their advancement to the clinic. Previous work has shown that the presence of pre-existing antiviral neutralising antibodies, which target the vector, can limit therapeutic efficacy by neutralising the vector before it has therapeutic effect^10,11^. Studies also demonstrated that HAdV-C5 could form high-affinity interactions with coagulation factor X, and/or platelets, resulting in their being trafficked to the liver, leading to degradation rather than delivery to the therapeutic target, resulting in lower efficacy^12–14^. This mechanistic understanding enabled the selection of low population seroprevalence adenoviruses (including ChAdOx1), and engineering of the capsid proteins to ablate many of these issues, as previously reviewed^15^.

Therefore, it is critical to fully investigate the vector-host interactions of ChAdOx1 on a mechanistic level. This will assist in understanding both how the vaccine generates immunity, and how it may lead to any rare adverse events, such as VITT. To this end, we determined the structure of the ChAdOx1 capsid and that of the tissue tropism determining fiber-knob protein. We provide the first robust evidence of ChAdOx1’s primary cell-binding receptor, demonstrating that the fiber-knob protein binds to coxsackie and adenovirus receptor (CAR) with high affinity, comparable to that of human adenovirus species C type 5 (HAdV-C5), while not interacting with membrane cofactor protein (CD46). Next, computational models suggest ChAdOx1 binds to PF4 via electrostatic interactions between PF4 and the hexon proteins, supporting recent observations by Greinacher *et al*^9^. Finally, we suggest a possible mechanism by which the association of ChAdOx1 and PF4 could result in VITT.

## Results

### The structure of the ChAdOx1/ChAd-Y25 viral capsid

Determined at 4.16Å resolution, ChAdOx1 possesses a prototypical icosahedral capsid structure (Fig.1A). As in other adenovirus structures solved by CryoEM, resolution is better on the more ordered interior of the capsid while the flexible components on the capsids’ exterior result in less well resolved signal (EFig.1). The asymmetric unit contains the expected penton monomer, one peripentonal, two secondary, and one tertiary hexon trimers, one peripentonal and one secondary copy of pVIII and one pIIIa protein (Fig.1B, C). We did not observe density for pVI or pVII.

**Figure 1:**
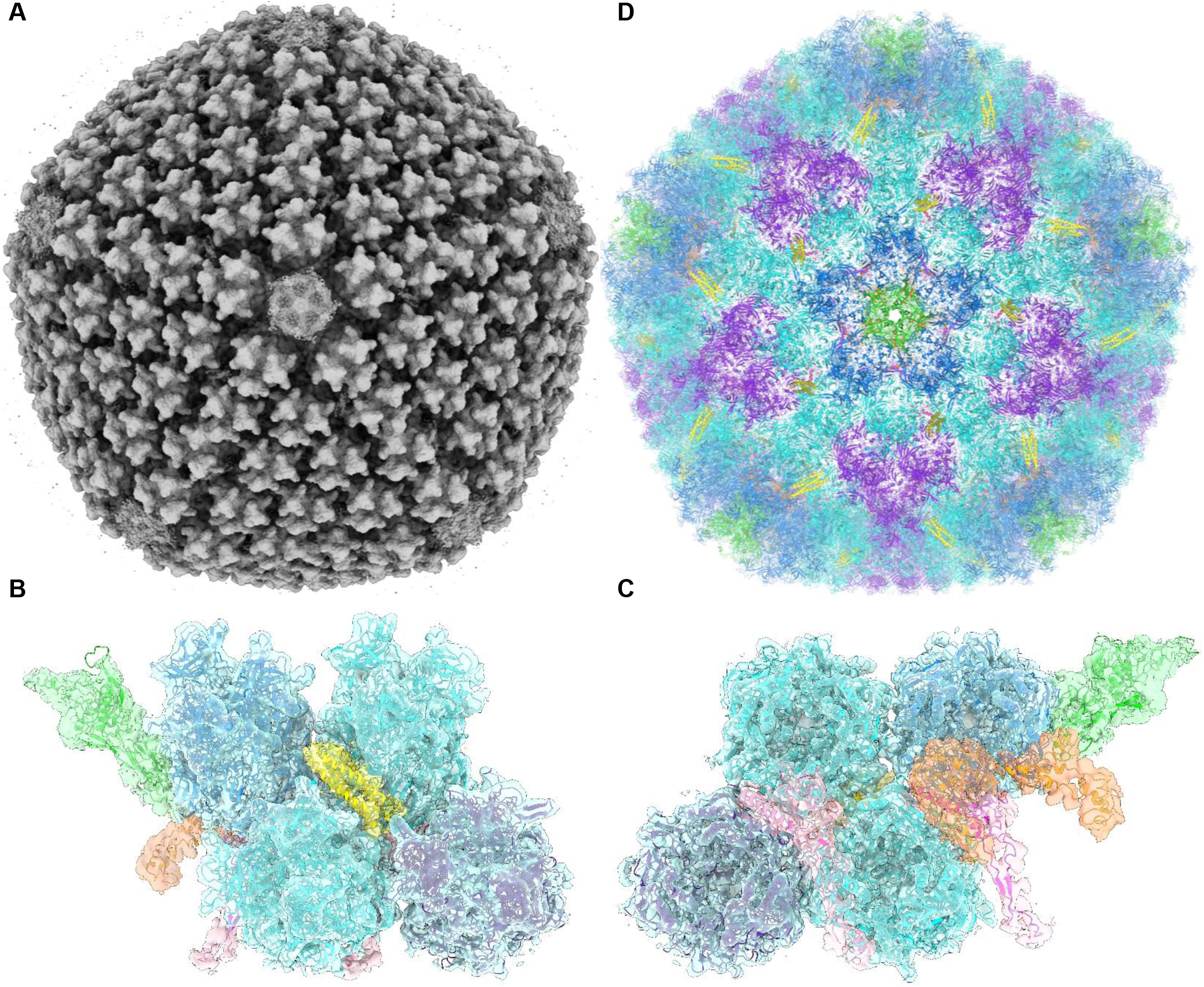
Overview of the ChAdOx1 viral vector capsid structure. Volume data for ChAdOx1 shows an archetypical adenovirus icosahedron (A) This volume resolved to show an asymmetric unit containing one penton copy (green), one trimeric peripentonal hexon (blue), two 2’ hexons (cyan), one 3’ hexon (purple), a 4-helix bundle corresponding to four copies of pIX (yellow), a peripentonal pVIII (magenta), a 2’ pVIII (pink), and a pIIIa protein (orange) seen from the capsid exterior (B) and interior (C) in their volume (grey). Repeating these asymmetric units with T25 icosahedral symmetry enables the reconstruction of a full ChAdOx1 capsid model (D).

The penton protein, forming a pentamer at each of the 12 icosahedral vertices, had relatively weak signal (Fig.1B, C), which may indicate a less well-ordered, or less stably interacting, protein than those observed in previous CryoEM structures of HAdV-D26 and HAdV-C5^16,17^. As in other reported structures the penton RGD loop (residues 305-335), which, in other adenoviruses is responsible for binding to integrins following attachment to the cell surface via the fiber-knob protein, was left un modelled due to a lack of signal, indicative of its flexibility.

pVIII (Fig.1C) was well resolved, barring residues 103-163, for which we observed no signal (Fig.1E), in line with the observations of other groups. Only partial density was observed for pIIIa, concentrated near the base of the penton, with weak signal occurring above the peripentonal pVIII and residues 313 to the N-terminus being unmodeled (Fig.1C). The hexon was well resolved, including the hypervariable regions (HVRs) on the exterior (Fig.1C), enabling a complete reconstruction. Reconstruction of the full capsid atomic model results in minimal clashes between asymmetric units and reconstructs a full icosahedral capsid (Fig.1D). The trimeric fiber protein, consisting of a long flexible shaft terminating in the globular knob domain, was not modelled as only short stubs of the shaft were visible due to its flexibility (EFig.1). Instead, the fiber-knob was solved by crystallography.

### CAR is a high affinity ChAdOx1 Fiber-knob receptor

Adenovirus fiber-knob protein is responsible for the primary virus-cell interaction during infection. To investigate primary ChAdOx1 receptors we solved the structure of the ChAdOx1 fiber-knob receptor. Data from one crystal were used for the final structure analysis, showing P1 symmetry with cell dimensions a=57.63Å, b=98.78Å, c=99.95Å, a=88.94°, b=86.29°, g=87.95°. Diffraction data were scaled and merged at 2.0Å resolution. The fiber-knob shows the expected three monomers assembling into a homotrimer with 3-fold symmetry, packed into an asymmetric unit containing 4 trimeric copies (EFig.2B). The electron density map was of sufficient quality to determine side chain conformations through the majority of the structure (EFig.2C, D). Super-positioning of the previously reported structure of HAdV-C5 fiber-knob and our structure of the ChAdOx1 fiber-knob protein show that, despite only 64.86% amino acid sequence homology, overall fold homology shows a root mean square deviation of 1.4Å (EFig.3A).

Using the existing structure of HAdV-D37 fiber-knob in complex with CAR we generated models of the HAdV-C5 (a known CAR interacting adenovirus), HAdV-B35 (a classical CD46 interacting adenovirus, which does not bind CAR), and ChAdOx1 fiber-knobs-CAR complex (EFig.3B-D). HAdV-C5 fiber-knob formed numerous polar contacts with CAR indicating a strong interaction (Fig.2A). HAdV-B35 fiber-knob formed just four polar contacts suggesting a weak interaction (Fig.2B), in line with previous observations^18^. ChAdOx1 fiber-knob also formed numerous polar contacts suggesting a strong interface similar to HAdV-C5 (Fig.2C). Interface energy calculations supported these observations, suggesting the strongest CAR interaction was formed by HAdV-C5, followed ChAdOx1, and then HAdV-B35 fiber-knobs (Fig.2D).

**Figure 2:**
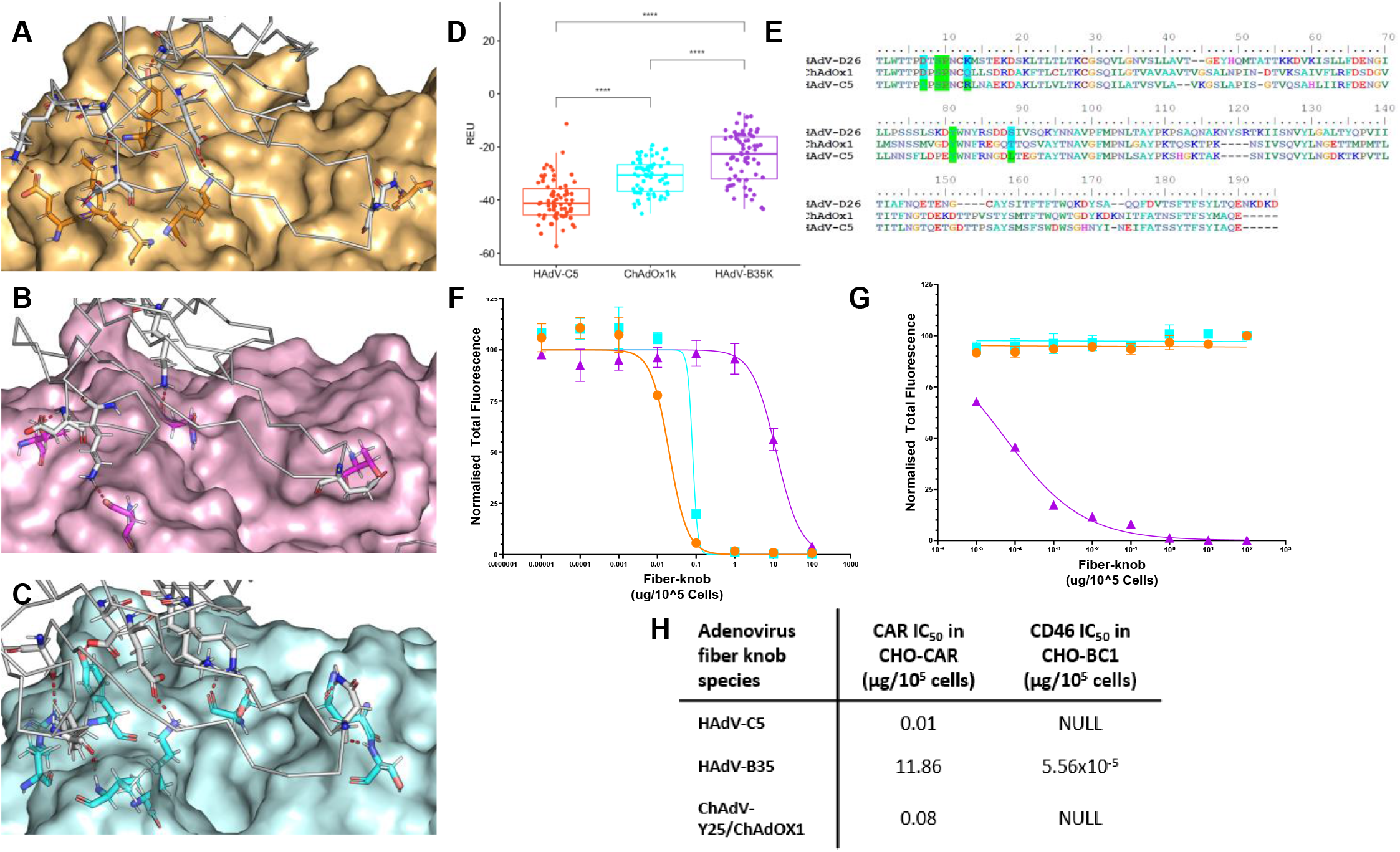
ChAdOx1 fiber-knob binds to CAR as a high affinity receptor. A model of HAdV-C5 fiber-knob in complex with CAR shows numerous contacts between CAR (grey) and HAdV-C5 (orange, A). A similar model of HAdV-B35 fiber-knob (purple) shows few contacts between long flexible sidechains (B) while the ChAdOx1 fiber-knob model (cyan) shows numerous contacts (C). Interface energy calculations performed support that HAdV-C5 is the strongest binder, followed by ChAdOx1, then HAdV-B35 (D), the sequence alignment of these fiber-knobs shows how HAdV-D26 and ChAdOx1 share many known CAR residues with HAdV-C5 (E). Antibody inhibition assays show HAdV-C5 is most able to block CAR binding in CHO-CAR cells, followed closely by ChAdOx1, and then distantly by HAdV-B35 (F), a similar assay in CHO-BC1 cells demonstrates that only HAdV-B35 fiber-knob is capable of binding to CD46 (G). The relative IC_50_’s of these assays are in given in the table (H).

Antibody inhibition assays in CHO-CAR cells, which express CAR but are negative for any other known primary adenovirus receptors, validated these findings biologically, assessing the ability of recombinant fiber-knob protein to block αCAR antibody binding. Consistent with the *in silico* findings, HAdV-C5 demonstrated the lowest IC_50_, indicating the strongest affinity for CAR, followed closely by ChAdOx1. HAdV-B35 demonstrated an IC_50_ three orders of magnitude higher than the others, likely representing non-specific inhibition resulting from the high protein concentration (Fig.2F) as it has previously been shown to be unable to utilise CAR as a cell entry receptor^18^. A similar analysis in CHO-BC1 cells, expressing CD46-BC1 isoform but no other known adenovirus receptors, determined that HAdV-B35, a known high affinity CD46 binding adenovirus, had a strong CD46 interaction, while HAdV-C5 (negative control) and ChAdOx1 fiber-knobs did not have a measurable ability to prevent αCD46 antibody binding, suggesting they do not bind (Fig.2G, H). Therefore, despite amino acid sequence differences, ChAdOx1 fiber-knob protein exhibits significant structural and functional homology to HAdV-C5.

### Apices of ChAdOx1 hexons have a strong electronegative potential

Adaptive Poisson-Boltzmann Solver (APBS) calculations performed upon models of the HAdV-C5, HAdV-D26, and ChAdOx1 hexon proteins, equilibrated by molecular dynamics, show that they are all electronegative in their apical regions (Fig.3A, B, C). ChAdOx1 is more electronegative than HAdV-D26 and HAdV-C5. Modelling a full facet of the icosahedral capsid of ChAdOx1 shows that several HVRs face into the space between hexons (Fig.3D) and are highly flexible creating a fluctuating amount of space between hexons (Fig.3E). Continuum electrostatic surface potential calculations across the facet of the ChAdOx1 icosahedron shows the trimeric apices of the hexons have a surface potential <-1.5k_B_T, creating an electronegative potential across most of the surface (Fig.4A). This charge is only interrupted in the crevices between hexons which are occupied by pIX helical bundles, where the surface potential rises to -0.5K_B_T (Fig.4A).

**Figure 3:**
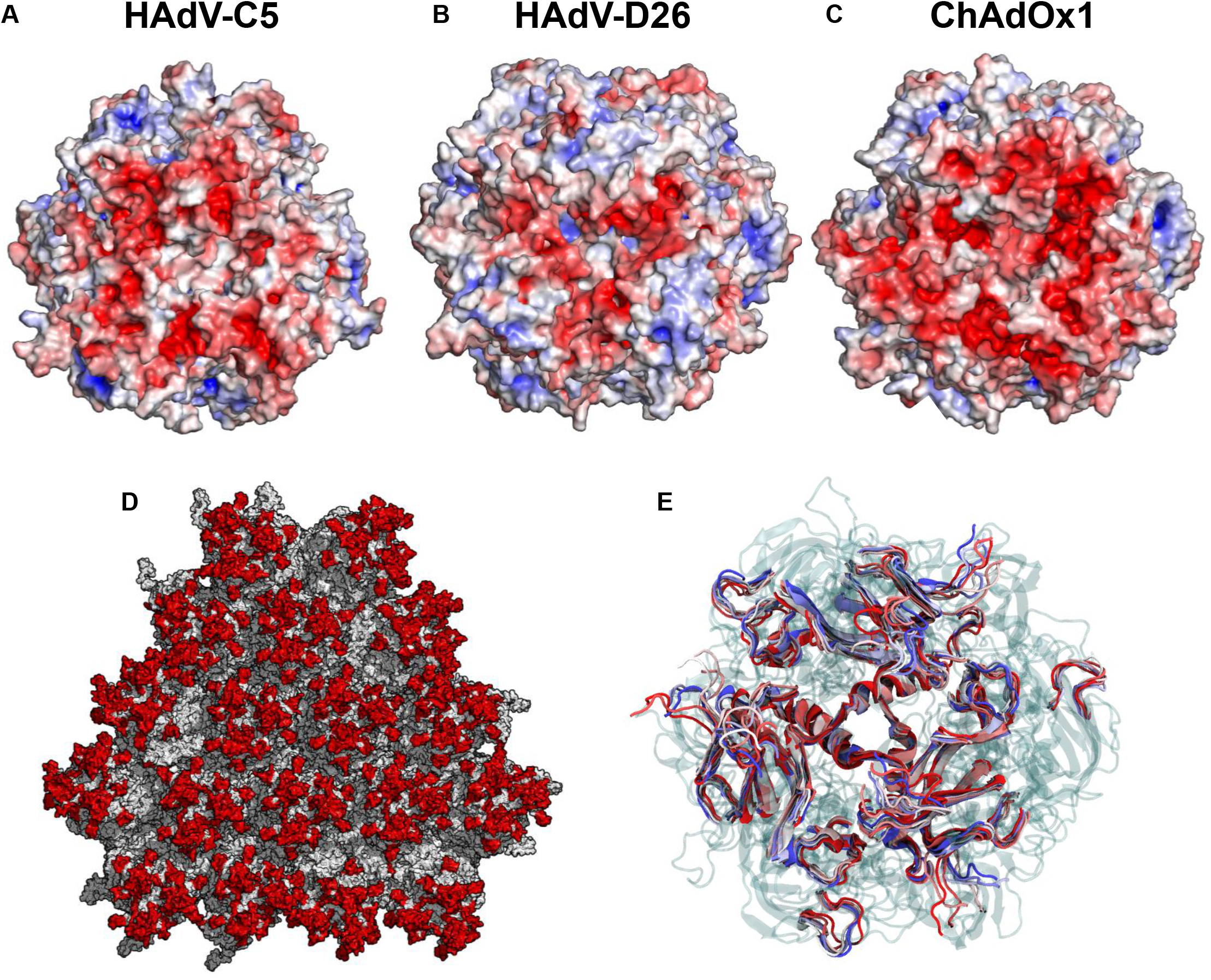
The ChAdOx1 hexon is strongly electronegative in its apex, and it’s flexible HVR’s contribute to this activity. HAdV-C5 is known to have an electronegative capsid surface contributed by its hexon protein (A), HAdV-D26 hexon is also electronegatively charged at its apex, but less so than HAdV-C5 (B). The hexon of ChAdOx1 is the most electronegative of the 3 proteins visualized (C) and part of this character is contributed by it’s HVR’s (red) which are seen to project into the spaces between hexons (D) and are highly flexible when assessed by molecular dynamics (E). APBS is visualize on a +/-5.0eV ramp from blue to red.

**Figure 4:**
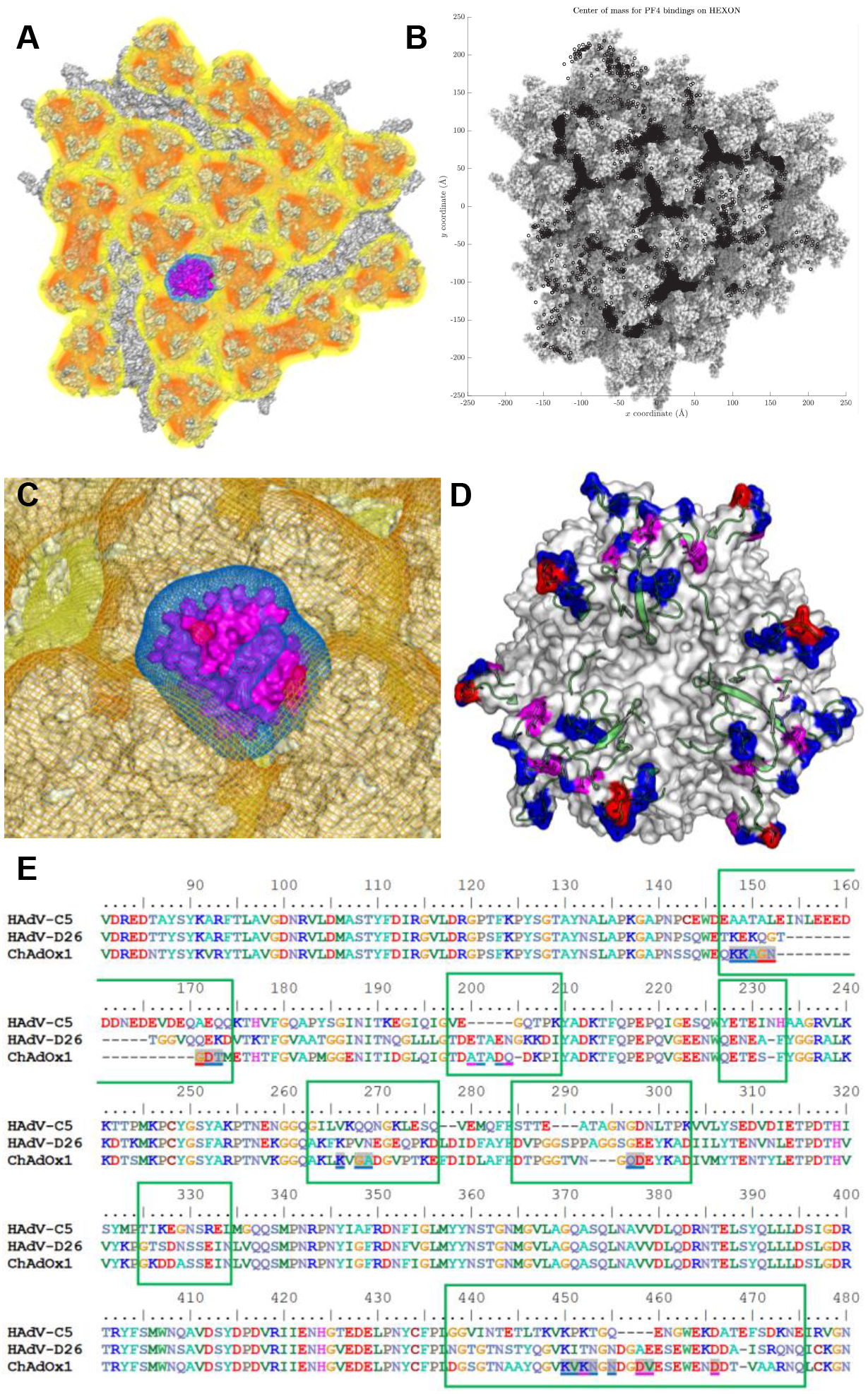
The ChAdOx1 capsid may form an electrostatic interaction with PF4. Visualizing the surface charge of ChAdOx1 at -0.5K_B_T (yellow), -1.0K_B_T (orange), and -1.5K_B_T (red) shows how the hexon apices are most positively charged and lead to a negative charge potential distributed over most of the icosahedral facet (A). Brownian dynamics PF4 in solution with the facet show the locations at which PF4 makes contact with the facet (black spots) over several thousand simulations (B) showing the most common interaction locus is the space between 3 hexons, where the PF4 is observed to sink into the space between hexons and turn its lateral, positively charged (Blue = +1.0K_B_T surface) face towards the oppositely charged hexons (C). Certain hexon residues are more commonly involved in the PF4 interaction (D), red residues interact >50% of the time, magenta >20%, blue 20-1%, with all these residues shown to be mounted upon HVR loops (green cartoon). The HVR containing hexon sequence is seen (E) with HVRs in green boxes and residues coloured according to the scheme in D.

### Brownian dynamics suggest an interaction between PF4 and ChAdOx1 capsid

We performed 400 independent 2μS Brownian dynamics simulations of PF4, a tetrameric electropositive protein (EFig.4A, B), with the ChAdOx1 facet. Results show freely diffusing PF4 frequently contacts the capsid surface between the hexons (Fig.4A, B, C). Closer observation of the interface at one of the most common interaction sites, the space between 3 hexon apices, shows PF4 penetrates between the hexons, and its positive charged surfaces face the oppositely charged hexons (Fig.3A, C). Analysis of the preferred orientation of PF4 when it contacts the hexons shows it is, usually, positioned to expose its electropositive faces towards the hexons electronegative HVRs, supporting an electrostatic mode of action (EFig.4C). A small number of hexon residues are seen to be involved in the majority of PF4 interactions (Fig.4D), all occurring in the HVRs (Fig.4E). The ChAdOx1 HVRs are highly flexible (Fig.3E) with root mean square fluctuations of 2Å (EFig.4D). ChAdOx1 and HAdV-D26 have similar flexibility (EFig.4D) but the former is more electronegative. This 2Å flexibility in the HVRs is expected to offer a favourable binding entropy which, together with its charge complementarity to PF4, is expected to facilitate ChAdOx1-PF4 recognition^19^.

## Discussion

ChAdOx1 shares the archetypal icosahedral, T=25, capsid common to adenoviruses. The fiber protein, which has 3-fold symmetry, protrudes from the pentons, which have 5-fold symmetry, at the vertices. This symmetry mismatch, combined with the inherent flexibility and propensity to break during concentration prior to grid preparation, has left them unresolved in our analysis. Limited information may be able to be obtained through local refinement methods, such as LocalRec^20^. It may be possible to achieve higher resolution from this data set by application of symmetry relaxation and expansion methods, however the high memory requirements of working with large high resolution particle images (1100pixels, 0.63 Å) have made this problematic on available hardware.

Though the overall fold of the ChAdOx1 capsid proteins is similar to other adenoviruses there are notable differences in character. ChAdOx1 hexon HVRs are different to other adenoviruses, as expected of a protein region which is, by definition, variable (Fig.3, 4). The ChAdOx1 HVRs are more similar in length and sequence to those of HAdV-D26 than HAdV-C5 (Fig.4E), displaying similar flexibility in molecular dynamics simulations (Fig.3E, EFig.4D). However, in terms of apical electrostatic surface potential, ChAdOx1 is more similar to HAdV-C5 than HAdV-D26, with APBS calculations performed on equilibrated hexon structures, showing that ChAdOx1 is most electronegative, closely followed by HAdV-C5, with HAdV-D26 being notably less electronegative (Fig.3A-C). This increased degree of negative surface potential may influence the strength of incidental charge-based interactions with other molecules.

We confirmed CAR as a high-affinity receptor for ChAdOx1 fiber-knob protein. Given the established ability of ChAdOx1 to infect human and chimpanzee cells, it follows that it uses a receptor which is conserved between these species for cell entry. CAR is an example of this, as Human and Chimpanzee CAR proteins have identical amino acid sequences (EFig.5)^1^. To the best of our knowledge, this is the first verified example of human and primate adenoviruses sharing a common adenovirus receptor protein. This is not a surprising result, as cross species utilisation of CAR has previously been observed. Both human and canine adenoviruses use CAR as a high-affinity cell entry receptor^21,22^. However, it is important to note that the formation of a productive infection is dependent upon more complex post-cell entry factors and a shared receptor does not equate to a shared pathogen.

While we have demonstrated that ChAdOx1 fiber-knob does not bind to CD46, another common adenovirus receptor, it does not preclude additional receptors. For example, HAdV-D37 and HAdV-D26 fiber-knob proteins have been shown to have dual affinity for CAR and sialic acid bearing glycans^23,24^. There could also, conceivably, be direct capsid-cell surface binding mechanisms, such as the hexon-CD46 interaction recently suggested for HAdV-D56, though it is unclear whether this reported low affinity interaction would be sufficient to form a productive infection *in vivo*^25^.

CAR has complex functions relating to junctional adhesion, and is predominantly expressed in epithelial tissues^26,27^. It has also been observed on the surface of human platelets, especially platelet aggregates^28^. It may, therefore, be tempting to try to link the affinity of ChAdOx1 for CAR with platelet aggregation and VITT. However, previous studies demonstrate that adenovirus-platelet aggregates are rapidly trafficked to the liver where they are sequestered by Kupffer cells and degraded^29^. While it was possible to induce thrombocytopenia in mice following an intravenous dose of 10^11^ replication deficient adenoviral particles^12^ (a dose which, by body weight, is ∼7,500X higher than that given, intramuscularly, in the ChAdOx1 nCoV-19 vaccine, assuming a 20g mouse, and a 75Kg human), this did not result in any thrombotic events. Rather, another study in Rhesus Macaques observed the opposite effect, longer clotting times, presumably as a result of the diminished platelet count and/or depletion of circulating coagulation factor X^30,31^. Therefore, we believe it is unlikely that direct association between ChAdOx1 and platelets contributes to thrombotic events, regardless of their CAR expression status.

A recent report on VITT by Greinacher *et al* suggested a direct association between PF4 and the ChAdOx1 capsid surface by transmission electron microscopy of ChAdOx1 nCoV-19 vaccine incubated with gold labelled PF4^32^. Our study presents a plausible mechanism for this interaction. Continuum electrostatic calculations and Brownian dynamics simulations suggest an incidental shape complementarity, stabilised by electrostatic interactions (Fig.3,4, EFig.4). This should be confirmed by detailed structural studies and does not preclude a specific protein binding site.

We observe that HAdV-D26 has similar capsid characteristics to ChAdOx1 but is less electronegative. Therefore, it seems plausible that HAdV-D26 could form a similar interaction with PF4, though the weaker electronegative potential may imply a weaker interaction.

VITT presents similarly to heparin-induced thrombotic thrombocytopenia (HITT), a disease where patients present with blood clots following administration of the thromboprophylactic drug heparin^33^. This disease appears to be driven by αPF4 auto-antibodies of sufficient affinity to cluster PF4 and create a multi-valent, presumably higher avidity, interaction between the antibody Fc-domains and FcγRIIa on the platelet surface, stimulating the platelet to activate and release further PF4. In the context of heparin, PF4 undergoes a conformational change facilitating the binding of, more common, lower affinity antibodies specific to the PF4-polyanion complex. This creates a positive feedback loop as antibodies bind to increasing copies of PF4, stimulating further platelet activation, culminating in the activation of the clotting cascade. This mechanism is described in detail in Nguyen *et al*^3^. To further summarise, this study suggests that whether αPF4 auto-antibodies induce thrombosis, or not, is a function of concentration and affinity to PF4.

Two studies, both from Greinacher *et al*, show that all patients who presented with VITT following ChAdOx1 vaccination (n=4, and n=28, respectively) tested positive for αPF4 antibodies^4^. Unlike those observed in HITT, αPF4 antibodies observed in VITT patients were predominantly of the sub-group which could bind to PF4 alone, rather than the PF4-heparin complex, as observed in HITT^32^.

We agree with the recent hypothesis from Greinacher *et al* that ChAdOx1 vaccination may be inducing the expression of high-affinity αPF4 antibodies, seeding aggregate formation and over-activation of platelets, resulting in VITT^9^. However, we propose a different mechanism by which this could be initiated. We suggest that the minor capillary injuries caused by the injection of the vaccine allow small quantities of ChAdOx1 to enter the blood. Once it has entered the blood it can associate with PF4, which will be secreted due to the immunogenic environment induced by the vaccine, by the mechanism described in Figure 4. At this point the ChAdOx1-PF4 complex may travel to the lymphatic system, either by diffusion or uptake and transport by monocytes, where they may stimulate pre-existing αPF4 memory B-cells to differentiate into plasma cells and secrete αPF4 antibodies, which generally takes 4-8 days^34^.

This differs from the proposal by Greinacher *et al*^9^ in the following ways. We do not propose EDTA as a contributing factor to VIT; while it is a component of the ChAdOx1 nCoV-19 vaccine formulation, it is not present in the Ad26.SARS2.S vaccine, which has also been observed to cause VITT^5,6,35^. We also do not propose a role for the HEK-293 cell contaminants observed in the vaccine, which Greinacher *et al* propose lead to systemic inflammation, release of PF4, and B-cell stimulation. Alternatively, we suggest trafficking of ChAdOx1 associated PF4 to the lymph is more likely to instigate memory B cell differentiation and the release of αPF4 auto-antibodies. We note that, VITT symptoms manifest 5-24 days following vaccination^9^. We suggest that if large immune complexes formed directly with vaccine constituents, it is more likely these would lead to platelet activation and clot formation immediately following vaccination, rather than >6 days following, when these constituents will likely have been cleared from the body. Following induction of high titre/high affinity αPF4 antibodies we are supportive of the mechanism suggested by Greinacher *et al*.

Key points remain to be addressed in exploring the mechanisms underpinning VITT. These include the association constant between ChAdOx1 (and other adenoviruses) and PF4, definitive evidence of whether VITT patients possess pre-existing PF4 specific B-cells, and if the adenovirus-PF4 complex can indeed result in their activation. It will also be useful to perform further analyses on patient data to identify the median time from vaccination to onset of VITT as a larger, more statistically significant, dataset becomes available.

Better understanding of the mechanism by which PF4 and adenoviruses associate presents an opportunity to engineer the capsid to ablate this interaction. ‘HVR swaps’ have been performed previously with the goal of reducing recognition of adenovirus vector by neutralising antibodies^36,37^. There is no reason that similar techniques could not be utilised to reduce surface electrostatic charge. Alternatively a more specific approach could be elected in which only key, electronegative, residues and those which may form key contacts with PF4 are removed or substituted. Therefore, modification of the ChAdOx1 hexon HVRs to reduce their electronegativity may solve two problems simultaneously: reduce the propensity to cause VITT to even lower levels, and reduce the levels of pre-existing immunity thus helping to maximise the opportunity to induce robust immune responses.

## Materials and Methods

### Propagation of ChAdOx1 virus

10 x T225 CellBind™ cell culture flasks (Corning) were seeded with approximately 5×10^6^ HEK-293 T-Rex cells (Invitrogen) each. Cells were cultured in DMEM (Gibco) supplemented with 10% heat inactivated fetal bovine serum and 1% penicillin/streptomycin (Gibco) until 80% confluent. Cells were infected with ChAdOx1.eGFP, provided by the Coughlan Lab (University of Maryland), at a multiplicity of infection of 0.01. Cells were then monitored for cytopathic effect (CPE). When culture media turned yellow media was replaced. Once CPE became evident, but the cells were not ready to be collected, yellowed media was supplemented with sodium bicarbonate buffer (pH7.4, Gibco) until it reddened. This was done to retain the virus released into the media. Once CPE was observed in >80% of the cell monolayer the cells (5-8 days post infection) were dissociated from the flask by knocking. The supernatant and cells were separated by centrifugation at 300g for 5mins, both were stored at -80C until ready for purification.

### ChAdOx1 Purification

ChAdOx1 containing media was clarified by centrifugation at 4000RPM for 10mins in a bench top centrifuge. Supernatant was then loaded into 38ml Ultraclear tubes compatible with the SW28 rotor (Beckman-Coulter) and centrifuged at 100,000g for 1hr in a Beckman-Coulter Optima XPN-100 ultracentrifuge. Supernatant was discarded and the pellet, which was slightly yellow and sticky, was resuspended in 5ml PBS (pH7.4, Gibco). This 5ml of PBS containing the ChAdOx1 from the supernatant was used to resuspend the infected HEK-293 T-Rex pellet. This solution was then mixed in a 1:1 ratio with tetrachloroethylene (TCE, Sigma-Aldrich) and shaken violently to ensure thorough mixing. The virus, PBS, TCE mixture was then centrifuged at 2000RPM in a benchtop centrifuge for 20minutes. The aqueous top layer of the solution was removed by pipetting and placed into a new tube. A second TCE extraction was then performed by adding a further 5ml of TCE to the aqueous layer, shaking, and centrifuging, as before. This is performed to ensure maximum removal cell debris. Previous purifications which excluded this second extraction showed fatty deposits when analysed by negative stain transmission electron microscopy and resulted in viral aggregation. Next, the top, aqueous, layer was extracted again and the remainder of the purification was performed using the 2 step CsCl gradient method, as previous described^38^, except for the following modification: during the final extraction of the virus containing band, which should have a crisp white appearance, the band was not removed using a needle through the side of the tube, but by pipetting. A P1000 pipette was used to slowly withdraw fluid from the tube, taking from the meniscus, careful not to disrupt the band. As much fluid was removed above the band as possible, then a new pipette tip was used to withdraw the band from the meniscus in as little volume as possible.

This band was loaded into a 0.5mL Slide-A-Lyzer cassette with a 100,000 MWCO (ThermoFisher) and dialysed against 1L of plunge freezing buffer (150mM NaCl, 1mM MgCl_2_, 20mM Tris, pH7.4). This solution was used to prepare grids for CryoEM.

### Cryo-EM Grid Preparation

UltrAuFoil grids (688-300-AU, Ted Pella Inc.) were glow discharged on a PELCO easiGlow (Ted Pella Inc.) for 30 seconds. A Vitribot Mark IV automated plunge freezer (ThermoFisher) was used for blotting and freezing grids in liquid ethane using the following protocol: the sample chamber was set to 25°C and 100% humidity, 2.5ul of sample was pipetted onto both sides of the grid, and a blot force of 1 was used for 3 seconds before each grid was plunged and transferred to cryo-boxes under liquid nitrogen.

### CryoEM Data collection

The vitrified specimen was images using an FEI Titan Krios transmission electron microscope (TEM, ThermoFisher) operating at an accelerating voltage of 300 keV with a Gatan K2 Summit direct electron detector (DED) camera (Pleasanton, CA) at a magnification of 37,33x, corresponding to a pixel size of 1.34Å/pixel at specimen level, 2875 movies, each consisting of 40 frames over the course of 8 seconds, were collected in super-resolution mode with a sub-pixel size of 0.67 Å^2^. Defocus targes cycled from -0.5 to -1.6μm and the dose rate was an average of 2.589 electrons/pixel/second with 1.1802 e-/Å2 per movie. Movies were saved in a 4-bit, LZW compressed TIF format.

### CryoEM Image Processing and Structure Determination

Datasets were processed using RELION 3.1.1^39^ on a workstation with an Intel I7-6800K 3.4GHz processor, 128Gb of memory, a 1Tb SSD drive, and 4 Nvidia GTX 1080 GPUs. At times when dynamic memory became limiting, swap space had to be increased to 600Gb to avoid errors. Movies were imported and motion correction was performed using RELION’s implementation of the MotionCorr2, with a binning factor of 2 and removing the first 3 frames. CTF estimation was performed using CTFFIND-4.1 executables within RELION. Approximately 8000 particles were picked manually. A particle diameter of 960Å was used to ensure well cantered particles. Particles were extracted with a box size of 1440 pixels and rescaled to 512 pixels. These particles underwent several cycles of 2D classification followed by subset selection to remove poor quality classes and regroup particles. An *Ab initio* model was generated without imposing any symmetry. This model was then aligned with its phase origin with the icosahedral symmetry using ‘relion_align_symmetry’ program before being used as a reference map for 3D classification. After further refining particle alignments by 3D classification the particles were re-extracted without rescaling and underwent further rounds of 2D class averaging isolate the best aligned classes for structure refinement. The remaining 4,300 particles were re-extracted with a box size of 1,100 to fit within the memory constrains of the workstation. This dataset underwent several rounds of 3D auto-refinement to determine optimal parameters and mask settings. Ultimately a mask excluding a central sphere of a radius of 285 Å was used to produce a post processed refined structure of 4.17 Å resolution as estimated by gold standard Fourier shell correlation^40^. At this step memory limitations required increasing the swap space size to finish the refinement. Bayesian polishing resulted in negligible improvements. Finally, Ewald’s sphere correction yielded a map with an average resolution of 4.16Å^41^.

### ChAdOx1 Capsid Model Building

The amino acid sequence of the ChAdOx1 virus capsid proteins are described in the genome sequence of the source chimpanzee adenovirus type Y25 (ChAdV-Y25) used to develop the ChAdOx1 vector (NC_017825.1). There are no differences between ChAdOx1 and Y25 in the capsid proteins.

The I-TASSER webserver^42^ or SWISS-MODEL^43^ was used to build homology models of the various ChAdOx1 capsid proteins. Initially, the atomic models obtained from the web-server were rigid body fitted to the capsid density map using ChimeraX^44^, first by manual placement and then using the fit in map algorithm, beginning with the hexons, then the penton, pVIII, pIX, pIIIa, respectively, until a full asymmetric unit had been assembled. After initial placement of the atomic model inside the density map, the model was fitted using the method molecular dynamics flexible fitting (MDFF) in explicit solvent^45,46^. To avoid fitting of atomic models into adjacent density, additional protein models were added surrounding the asymmetric unit to occupy those regions of the density map. This ‘buffered’ asymmetric unit was then subjected to another round of MDFF in explicit solvent to improve the quality of fit to the density map, while accounting for protein-protein interactions between protein sub-units. Models were then inspected and loop regions with low-resolution density were manually modelled using ISOLDE^47^. The model was further refined in MDFF to address clashes and further improve the local fits. All MDFF simulations were performed in NAMD 2.14^48^ using the CHARMM36^49^ force field for protein, water and ions at 300 K. The initial system for every MDFF simulation was prepared using the software Visual Molecular Dynamics, VMD 1.9.3^50^. The potential energy function (U_EM_), obtained by converting from the EM density map was applied to the protein backbone during flexible fitting with a g-scale of 0.3. In addition, restraints were applied to maintain secondary structure, cis-peptide and chirality in protein structures, during flexible fitting. Finally, residues for which there was no signal were deleted from the model.

### Crystallization and structure determination of the ChAdOx1 Fiber-knob protein

#### Production and Purification of ChAdOx1 fiber-knob protein

The method used to purify ChAdOx1 fiber-knob protein is identical to that described for HAdV-D26 and HAdV-D48, previously. To summarise, SG13009 E.coli containing the pREP-4 plasmid were transfected with pQE-30 expression vector containing the ChAdOx1 fiber-knob transgene, located C-terminal to the 6His tag site, consisting of the 13 residues proceeding the TLW motif to the terminal residue. These *E*.*coli* were selected in 100μg/mL of Ampicillin and 50μg/mL of Kanamycin. Were cultured in 25ml LB broth with 100µg/ml ampicillin and 50µg/ml kanamycin overnight from glycerol stocks. 2L of Terrific Broth (Terrific broth, modified, Sigma-Aldrich) containing 100µg/ml ampicillin and 50µg/ml kanamycin were inoculated with the overnight culture and incubated at 37°C for 4 hrs until it reached an optical density (OD) of 0.6 at λ570nm. Once it reached the OD0.6 IPTG at a final concentration of 0.5mM was added and the culture was incubated for 18hrs at 21°C. Cells were harvested by centrifugation at 3,000xg for 10 mins at 4°C, resuspended in lysis buffer (50mM Tris, pH8.0, 300mM NaCl, 1% (v/v) NP40, 1mg/ml Lysozyme, 1mM β-Mercaptoethanol, 10mM Imidazole), and incubated for 15min shaking at room temperature. Lysate was then centrifuged at 30,000xg for 20 min at 4°C and filtrated through 0.22µm syringe filter (Milipore, Abingdon, UK). The filtered lysate was then loaded to 5ml HisTrap FF nickel affinity chromatography column (GE Life Science) at 2.0ml/min and washed with 15ml into elution buffer A (50mM Tris-HCl or- Base pH8.0, 300mM NaCl, 1mM β-Mercaptoethanol). Elution was done using a gradient rate of 20min/ml from of buffer B (50mM Tris pH8.0, 300mM NaCl, 1mM β-Mercaptoethanol, 400mM imidazole). Collected fractions were concentrated by centrifugation using a Vivaspin 10,000 MWCO column (Sartorious, Goettingen, Germany). and analyzed by a SDS-PAGE gel stained with Coomassie Blue (correct bands are approximately 25kDa). A second round of purification was performed using a GE 10/300 GL Increase Superdex 200 (GE Life Science) size exclusion chromatography at 0.5 ml/min and washed with 50ml of Baker Buffer (50mM NaCl, 10mM Tris pH 7.6). Fractions were then analysed using an SDS-PAGE gel stained with Coomassie Blue to check purity and molecular weight.

#### Crystallization conditions

Final protein concentration was 13.5 mg/ml. For crystallization The BCS and PACT Premier commercial crystallization screens (Molecular Dimensions) were used. Crystals were grown at 20°C in sitting drops containing 1:1 (v/v) ratio of protein to mother liquor. Microseeding was required for optimal crystal growth. Microseeding experiment was set-up using Mosquito crystallization robot. Crystals appeared between 5-14 days.

Crystallization condition for solved structure was 0.1 M Calcium chloride dihydrate, 0.1 M Magnesium chloride hexahydrate, 0.1 M PIPES pH7.0, 22.5 % v/v PEG Smear Medium and 0.2 M Calcium chloride dihydrate, 0.1 M Tris pH8.0, 20 % w/v PEG 6000.

#### Structure determination

Diffraction data were recorded on DIAMOND beamline DLS-I03, using GDA to control data collection. Automatic data reduction was completed with XDS and DIALS, and equivalents scaled and merged with AIMLESS and TRUNCATE^51^. The unique data set was used in PHASER to solve the structure with Molecular Replacement, using a search model prepared by SWISS-MODEL based on the structure of Adenovirus C5 fiber knob protein, PDB entry 1KNB^52^. Repeated cycles of graphics sessions in COOT^53^ and refinement in REFMAC5 resulted in the final model presented in this manuscript. It became clear that the data set suffered from twinning, which was automatically determined in REFMAC5 as 1 twin law, with a fraction of ∼0.15. Details of data collection and refinement statistics are included in Extended figure 2A.

### Modelling of fiber-knob CAR interfaces

The pre-existing structure of the HAdV-D37 fiber-knob in complex with CAR D1 (PDB 2J12)^21^ was used as a template by which to fit other fiber-knob proteins, as previously described. The pre-existing structure of HAdV-C5 (PDB 1KNB)^52^ and HAdV-B35 (PDB 2QLK)^54^ were aligned to the HAdV-D37 fiber knob in PyMOL using the ‘cealign’ command, as was the fiber-knob structure of ChAdOx1^55^. New models were saved containing the three CAR chains and one of the fitted chimeric fiber-knob trimers. These homology models underwent 10,000 steps of energy minimization using a conjugate gradient and line search algorithm native to NAMD^48^ and equilibrated by a short 2ns molecular dynamics simulation. These models, seen in figure 2, A-C and extended figure 3, were imaged and binding interaction between the all three fiber-knob and CAR protein-protein interfaces were scored at each frame of the MD trajectory using the Rosetta InterfaceAnalyzer tool^56,57^.

### Determination of relative IC_50_ values of CAR and CD46 binding for Fiber-knob proteins

Antibody binding inhibition assays were performed as previously described^18^. CHO-CAR and CHO-BC1 cells were harvested and 40,000 cells per well were transferred to a 96-well V-bottomed plate (NuncTM; 249662). Cells were washed with cold PBS prior to seeding and kept on ice. Serial dilutions of recombinant soluble knob protein were made up in serum-free RPMI-1640 to give a final concentration range of 0.00001–100 μg/10^5^ cells. Recombinant fiber knob protein dilutions were added in triplicate to the cells and incubated on ice for 30 minutes. Unbound fiber knob protein was removed by washing twice in cold PBS and primary CAR RmcB (Millipore; 05-644) or primary CD46 MEM-258 (ThermoFisher; MA1-82140) antibody was added to bind the appropriate receptor. Primary antibody was removed after 1 h incubation on ice and cells were washed twice further in PBS and incubated on ice for 30 minutes with Alexa-488 labelled goat anti-mouse secondary antibody (ThermoFisher; A-11001). Antibodies were diluted to a concentration of 2 μg/mL in PBS. Cells were washed and fixed using 4% paraformaldehyde and staining detected using BD Accuri™ C6 cytometer (BD Bioscience). Analysis was performed using FlowJo v10 (FlowJo, LLC) by sequential gating on cell population, singlets and Alexa-488 positive cells compared to an unstained control. Total fluorescence intensity (TFI) was determined as the Alexa-488 positive single cell population multiplied by median fluorescence intensity (MFI) and IC_50_ curves were fitted by non-linear regression using GraphPad Prism software to determine the IC_50_ concentrations.

### Sequence Alignments

Sequence alignments were performed using the Clustal Omega algorithm as implemented in Expasy^58^.

### Electrostatic surface calculations

The electrostatic effect from the icosahedral facet of ChAdOx1 on any point **r** = (*x, y, z*) of its surrounding environment is quantified by its electrostatic potential at **r**, which is denoted as *V*(**r**). This potential *V* was determined using the code, Adaptative Poisson-Boltzmann Solver (APBS)^59^. APBS computed *V* using the charge distribution of ChAdOx1, the ion concentration of the environment surrounding ChAdOx1, and the permittivity of the environment. The computed *V* thus took into account the presence of counter ions in and the polarizability of the environment, which was a bulk of water molecules in our study. As the sign, + or -, of *V* (**r**) is highly correlated with the number of positive or negative charges around **r**, the potential for platelet factor 4 (PF4) was also computed to visualize the overall charge distribution of PF4 upon its binding to ChAdOx1.

### Brownian dynamics simulation of the icosahedral facet in solution with PF4

Multi-replica Brownian dynamics (BD) simulations were performed using the code, Atomic Resolution Brownian Dynamics (ARBD)^60^. During the simulations, copies of PF4 were treated as rigid bodies diffusing in the neighbourhood of the icosahedral facet of ChAdOx1 in solution.

The equation of motion (EOM) obeyed by each copy of PF4 for its diffusion in BD simulations is the over-damped Langevin dynamics. This EOM requires knowledge of the forces from the environment on PF4 as well as its damping coefficients with the environment. These coefficients, translational damping coefficients and rotational damping coefficients, were determined using the code, Hydropro^61^.

The forces from the environment on PF4 consisted of 2 parts. The 1^st^ part consisted of the random forces from thermal fluctuations. These random forces were automatically generated by ARBD during simulations based on the system temperature *T*, which is 310K here. The 2^nd^ part consisted of the electrostatic force and the Van der Waals (VdW) force on PF4 from the icosahedral facet. The electrostatic force on PF4 with its centre of mass (COM) at any point **r** was computed by ARBD automatically using the electrostatic potential *V* of the icosahedral facet and the charge distributions of PF4 around **r**. Similarly, we specify the VdW force between the icosahedral facet and PF4 using a potential, the Lennard-Jones (LJ) potential, denoted as *U* here. This potential *U* was computed using the code, Visual Molecular Dynamics (VMD)^50^. LJ parameters needed for this computation were adopted from the Charmm36m force field^62^.

The various potentials for the icosahedral facet and the damping coefficients for PF4 were feed into ARBD for multi-replica BD simulations. Our work employed 16 replicas of simulations in parallel. Each replica lasted for a simulation time of 2μs and simulated 25 copies of PF4 diffusing around the icosahedral facet. During the simulations, these 25 copies of PF4 were only allowed to interact with the icosahedral facet and did not interact with one another. Thus, together, we performed 400 independent diffusion simulations with PF4. During each of these simulations, PF4 was allowed to bind to, or unbind, from the icosahedral facet. It is important to note that the residence time for each of these binding events is usually 2-3 order of magnitudes different compared with the real binding time. This is because the post-binding conformational changes of PF4, which often stabilize the interaction, are not sampled. Further, the simulations are performed in an implicit solvent environment, which is known to accelerate protein dynamics and diffusion, even in MD.

## Acknowledgments

We acknowledge Diamond Light Source for time on DLS-I03 under proposal mx-20147, and NSF grant 1531991 for support for the Titan Krios at ASU’s Eyring Materials Center at Arizona State University.

We would like to acknowledge Amy S. Codd in the Vaccine and Infectious Disease Division at the Fred Hutchinson Cancer Research Center, Seattle, USA, for her important insights into the immunological mechanisms discussed in this study and review of the manuscript. Finally, we wish to thank Lynda Coughlan in the School of Medicine, University of Maryland, Baltimore, USA, for important revisions to the manuscript and discussions regarding adenovirus vaccinology.

---

**Extended Figure 1:**
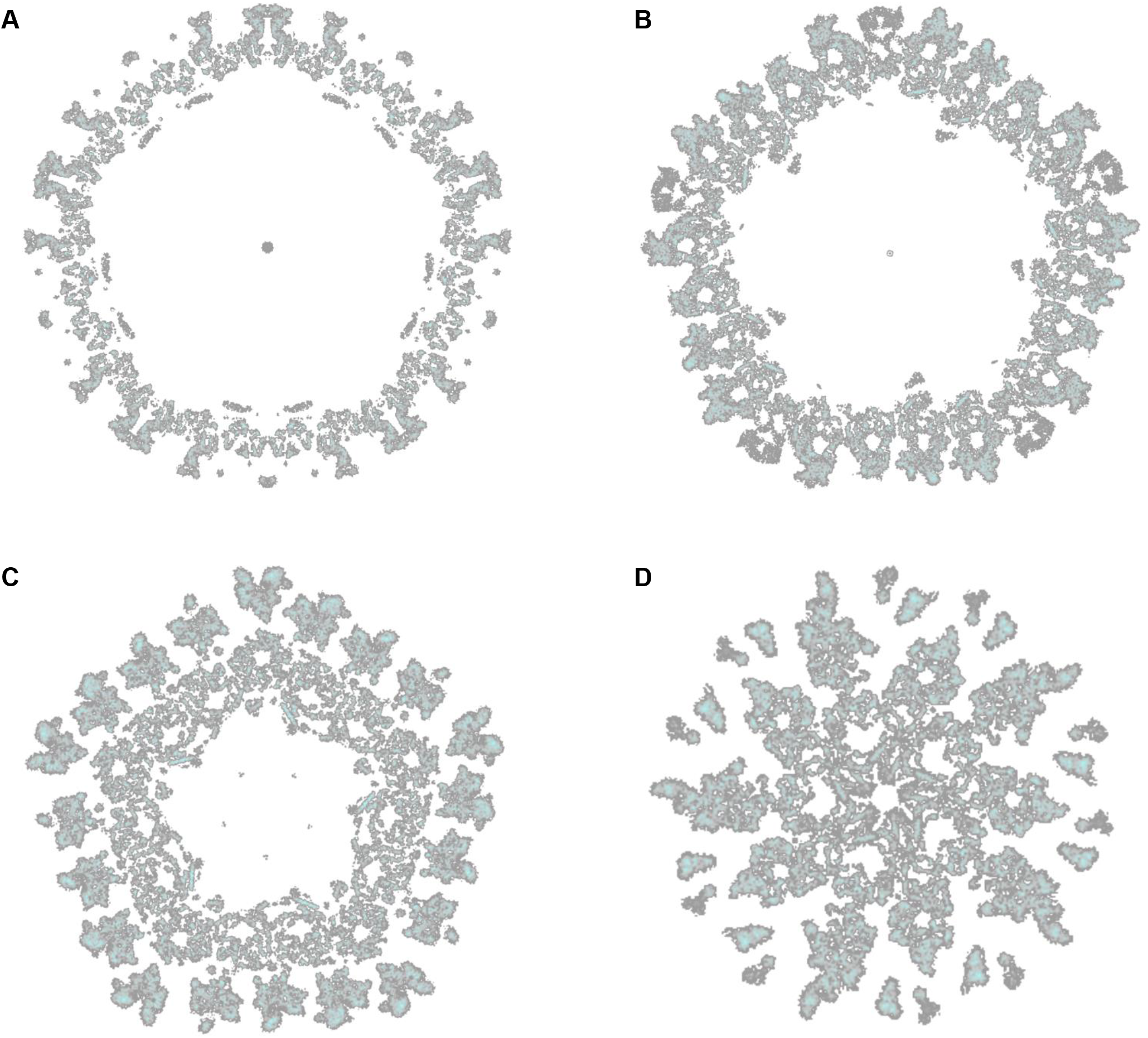
Slices through the ChAdOx1 CryoEM volume show the capsid interior has higher resolution information than the exterior which contains more flexible regions. An equatorial slice (A) shows greater detail on the capsid interior, revealed in more detail by slices at points further along the 5-fold axis (B, C, D).

**Extended Figure 2:**
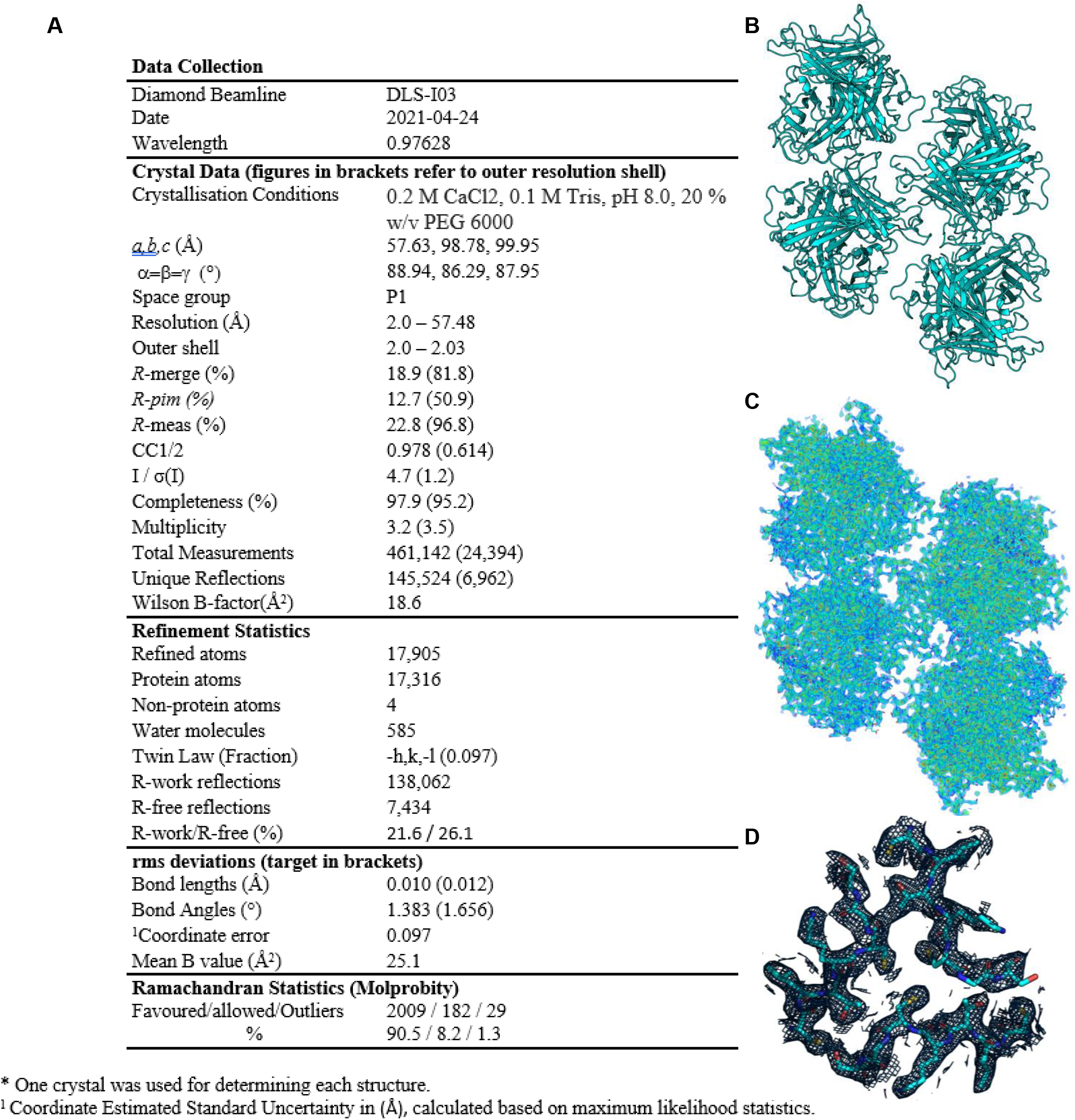
Crystalisation of ChAdOx1 fiber-knob protein results in 4 copies of the expected trimer per asymmetric unit and reveals side-chain locations. Acceptable refinement statistics were achieved for the fiber-knob protein of ChAdOx1 (A). The crystal structure was solved with 12 copies of the monomer in the asymmetric unit, packing to form 3 trimeric biological assemblies (B). Density was sufficient to provide a complete structure in all copies (C, volume rendered in 0.5σ steps from red, 3.0σ, to dark blue), and was able to resolve side chain orientations reliably throughout the core fold (D, mesh shown at σ=1.0).

**Extended Figure 3:**
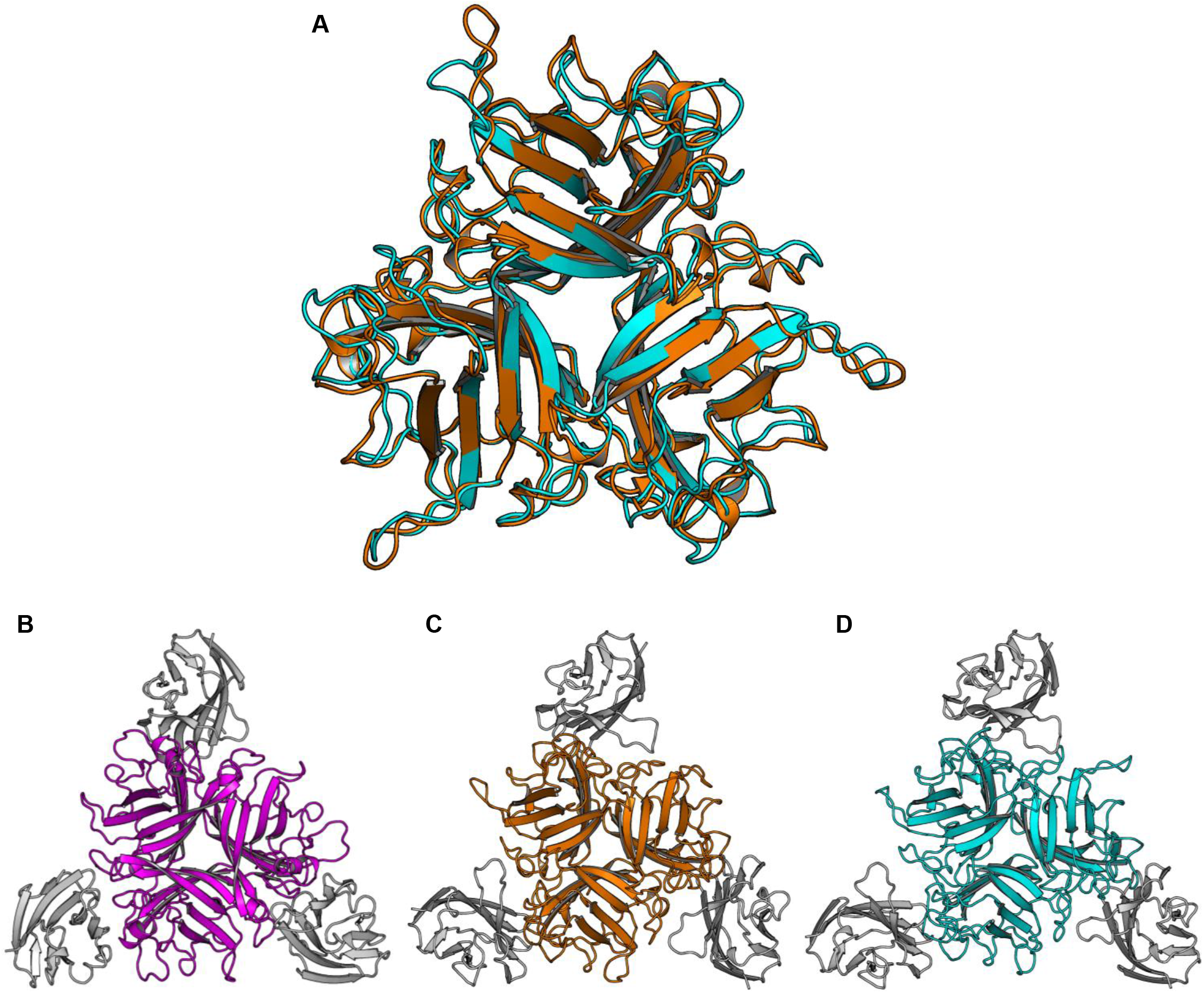
ChAdOx1 has homology to species C adenovirus 5, and models suggest binding to CAR. The crystal structure of the ChAdOx1 fiber-knob protein (cyan) exhibits a similar fold to that of HAdV-C5 (orange, A). Using PDB 2J12 as a template the fiber-knob structures of HAdV-B35 (Purple), HAdV-C5, and ChAdOx1 were aligned with CAR in a potential binding pose and equilibrated by molecular dynamics (B, C, D).

**Extended Figure 4:**
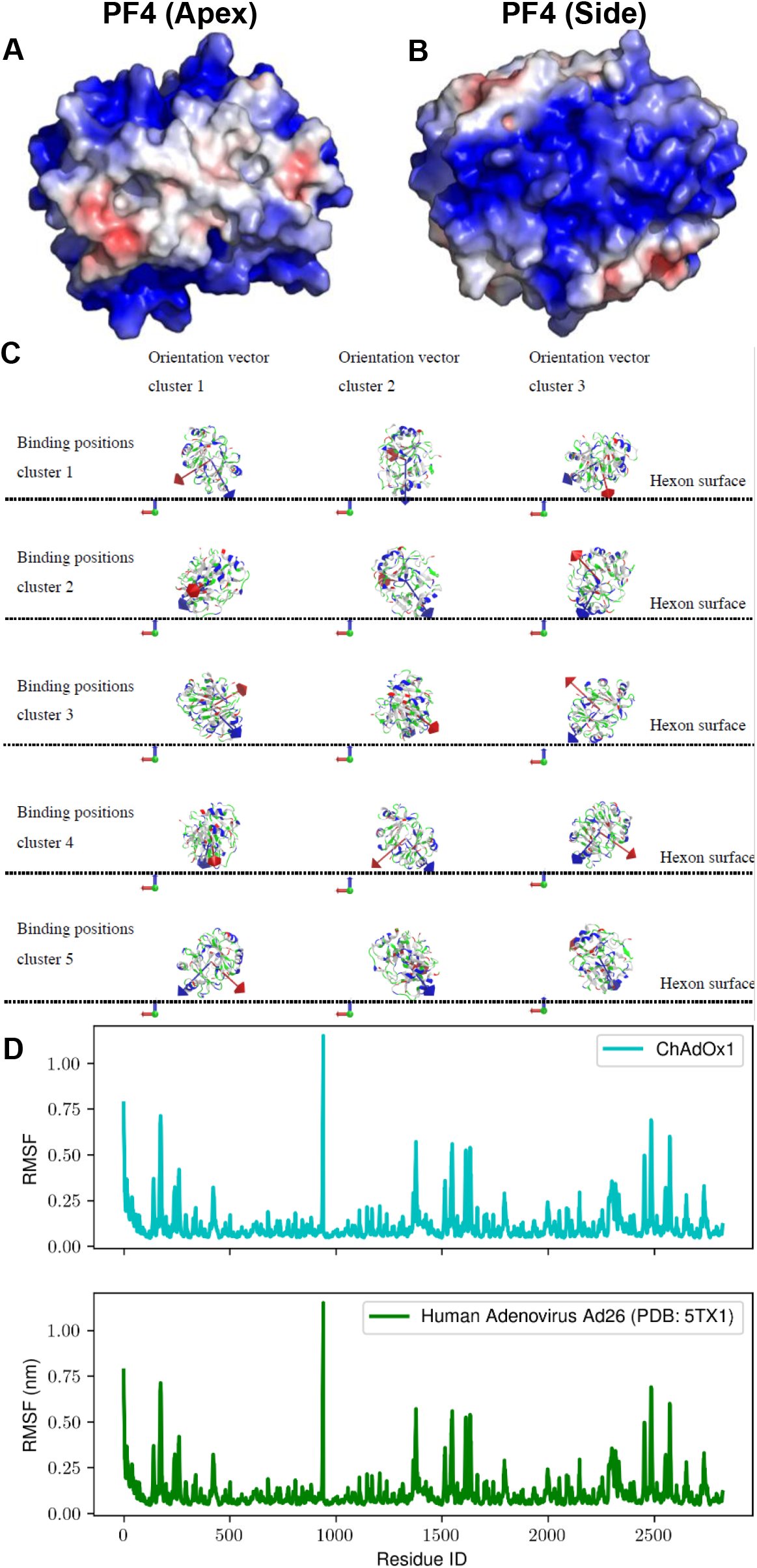
The electronegative surfaces of PF4 are oriented towards the electropositive faces of the hexons, which have flexible HVRs, and may facilitate their interaction. PF4 is an exceptionally electronegative protein around its longitudinal faces (A,B), and when it forms interactions with the ChAdOx1 capsid surface it usually positions itself such that its charged faces are oriented to face the hexon HVRs (C), the plane of the hexons is shown as a sashed line. The HVRs are determined to be highly flexible in both ChAdOx1 and HAdV-D26, by molecular dynamics (D). APBS is visualized on a +/-5.0eV ramp from blue to red.

**Extended Figure 6:**
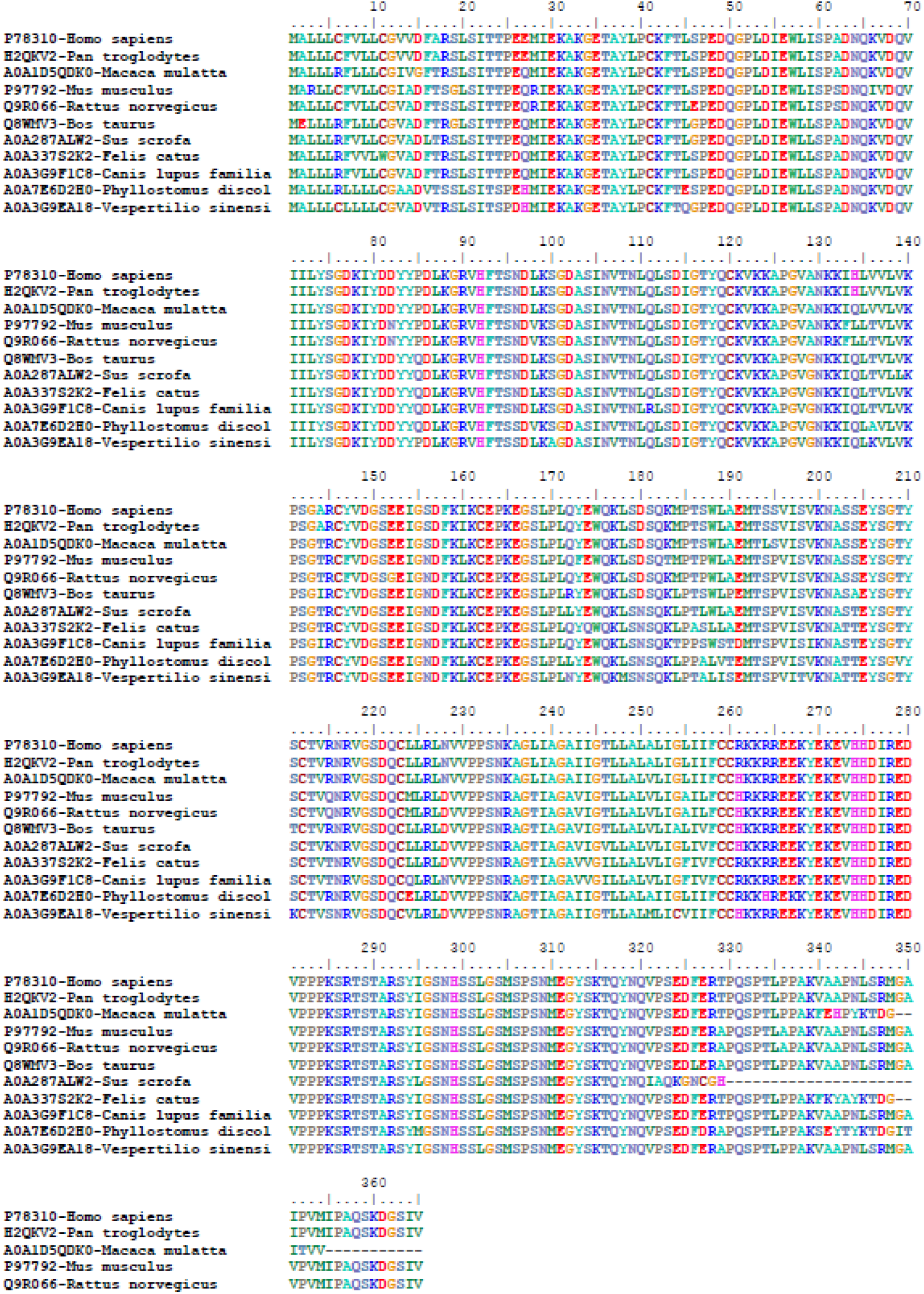
CAR is a highly conserved protein across a range of scientifically important, domestic, and agriculturally significant species. Humans (homo sapiens) and Chimpanzees (Pan Troglodytes) share a 100% sequence identity for their canonical CAR isoform. Sequences in this alignment taken from the indicted UniProt accession numbers.

